# Cell-free prediction of protein expression costs for growing cells

**DOI:** 10.1101/172627

**Authors:** Olivier Borkowski, Carlos Bricio, Michaela Murgiano, Guy-Bart Stan, Tom Ellis

## Abstract

Translating heterologous proteins places significant burden on host cells, consuming expression resources leading to slower cell growth and productivity. Yet predicting the cost of protein production for any gene is a major challenge, as multiple processes and factors determine translation efficiency. Here, to enable prediction of the cost of gene expression in bacteria, we describe a standard cell-free lysate assay that determines the relationship between *in vivo* and cell-free measurements and γ, a relative measure of the resource consumption when a given protein is expressed. When combined with a computational model of translation, this enables prediction of the *in vivo* burden placed on growing *E. coli* cells for a variety of proteins of different functions and lengths. Using this approach, we can predict the burden of expressing multigene operons of different designs and differentiate between the fraction of burden related to gene expression compared to action of a metabolic pathway.

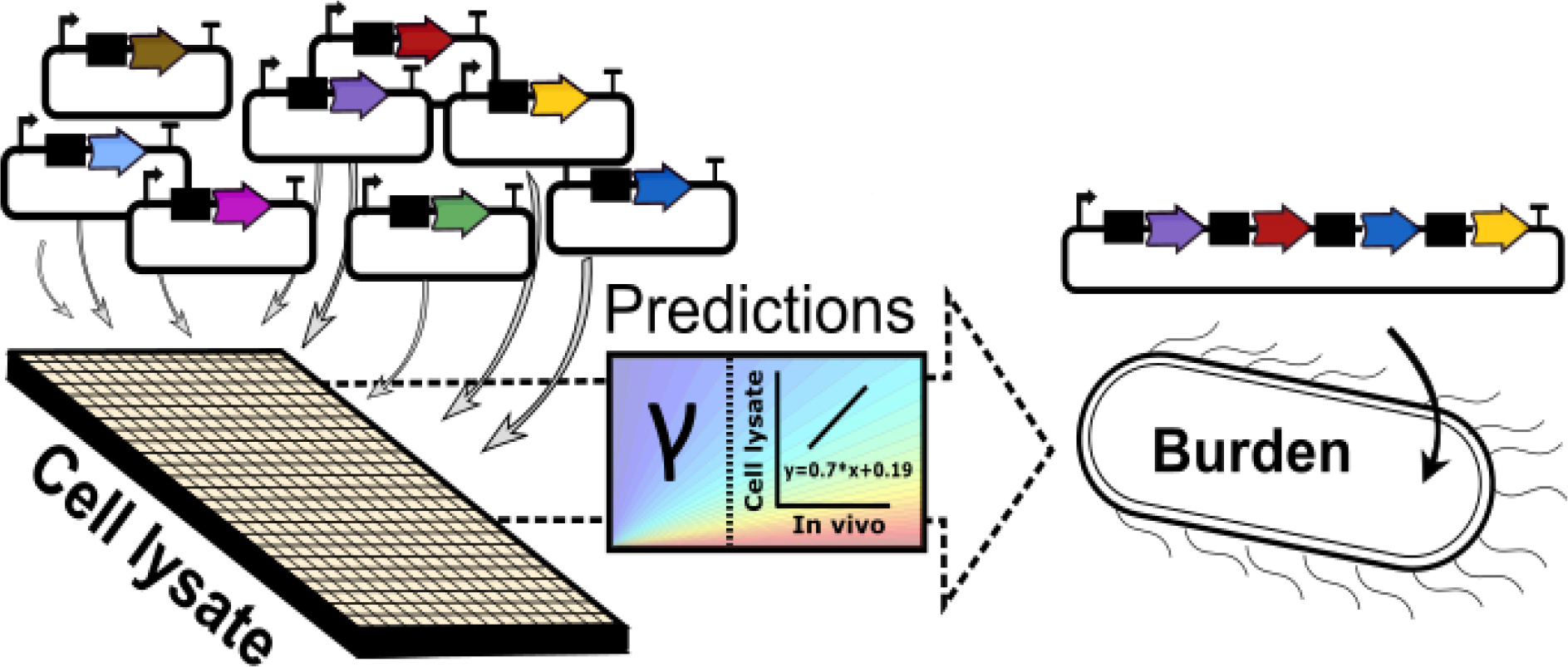

## Introduction

To be able to build systems with increasingly more genes, not only is precise gene expression desirable, but it is also essential to have an understanding of the burden these will place on the host cell so that designs can be optimised to ensure robust growth and to prevent the deleterious mutations that arise in high-burden systems ^1,2,3^. For any given gene, its burden is in the first instance the resource cost of maintaining and expressing the gene as needed ^4,5,6,7^. If the gene encodes a function, for example an enzyme, the impact of this can further cause a more specific role-based metabolic burden that adds to the expression burden, e.g. by consuming host cell metabolites and co-factors ^8,9^. Research primarily in the model bacteria *E. coli*, has demonstrated that a lack of understanding of the burden of expressing additional genes affects our ability to predictively engineer cells ^10,11,12,13^.

To predict the burden of expressing a new gene from a synthetic construct, we first need to understand how much it is expressed and how this affects the cell’s capacity for its own gene expression by consuming its resources. With synthetic biology tools and software it is now possible to define at the DNA sequence level both the amount of transcription (via the promoter) and the rate of translation initiation (via the RBS sequence) ^14,15^. Yet, the efficiency of translation is also known to be dependent on nucleotide composition ^16^, secondary structure of the mRNA ^17^, translational pausing ^18^, the presence of rare codons or the use of rare amino acids ^19,20^, or in most cases combinations of all of the above and more. Given this complexity it is not surprising that it is currently impossible to look at a given gene DNA sequence or protein amino acid sequence and predict what impact its expression will have on the host cell.

*In vivo* measurements are therefore still essential for predicting the cost of protein expression. The relative costs of translation of proteins can be inferred from the *in vivo* outcomes of competition for resources, either simply by measuring changes in growth rate measurements ^21^ or by quantifying decreased production rate of a “capacity monitor” gene; a constitutively expressed and chromosomally-integrated reporter that acts as a proxy of available expression resources in the cell ^4^. However, *in vivo* measurements are a low-throughput process requiring time-consuming cloning and growth experiments ^22,23^. They often generate hard-to-deconvolute results because burden slows growth and potentially triggers secondary changes in global gene expression behaviour.

Cell-free lysates are a new emerging tool in synthetic biology that represents a promising approach for quantifying burden as they constitute simpler, non-growing genetic expression systems akin to cells, that effectively capture a snapshot of the same gene expression machinery. Indeed, the expression of protein-encoding constructs using *E. coli* cell lysates has been shown to match *in vivo* performance in many cases ^24,25,26^. It also allows users to rapidly characterize a large number of synthetic constructs in parallel ^27^, and batches of cell lysates can be prepared and stored frozen, purchased from companies or can be customised in terms of their make-up ^28^.

Here, we combined cell lysate experiments, *in vivo* measurements in corresponding *E. coli* cells and mathematical modelling to demonstrate prediction of the impact of different genes and multigene systems on growing *E. coli* cells. We measured the resource consumption of a variety of protein coding sequences in both cell lysates and *in vivo*. Through this, we show that the competition for resources between a capacity monitor construct and the different protein coding sequences showed a correlation between cell lysate and *in vivo* data. Using a ribosome flow model, we extend predictions of burden in cells to systems expressing multiple genes of interest with different translation efficiencies, and in different growth conditions. By combining these efforts, we provide a novel method for rapid *in vitro* screening of synthetic parts designed to enable the prediction of the behaviour and expression efficiency of synthetic constructs and the impact they have on their host cell.

## Results

### Cell-free protein synthesis using *E. coli* lysates provides a rapid platform to quantify resource competition

Current methods to quantify burden all rely on *in vivo* measurement of cell performance and growth when hosting and expressing synthetic constructs ^4,29,30^. As such, it typically requires several days of cloning, verification and assaying to obtain data that can be used to determine a gene’s burden. However, groups using ‘cell-free’ protein synthesis systems have recently shown that the design-build-test cycle used in synthetic biology can be accelerated by characterising expression from DNA directly added to *E. coli* lysates ^24,25,26,27^.

*In vivo*, the burden of expression of synthetic constructs is caused by competition between the added genes and native genes for the resources needed for gene expression. Previously, we have shown that this can be measured by a decrease in the capacity of a cell to express a standard measurable gene chromosomally-integrated into *E. coli* that acts as a “capacity monitor” ^4^ (**Figure 1A**). Similarly, in cell-free experiments it has also been possible to observe competition for gene expression resources when using two different plasmids within the same cell lysate mix ^31^. Therefore, as a proxy to measure resource competition in cell lysate, we constructed and tested a low-copy plasmid-based version of our previous capacity monitor (**Figure 1B**) where *superfolder gfp* is expressed constitutively from a synthetic promoter with a strong RBS. Using the maximum GFP production rate (max dGFP/dt, see **Supplementary Figure 1A**) measured during characterisation with this plasmid in cell lysates, we sought to measure available expression capacity and identify resource competition. To do this we first simply measured the max GFP production rate in 10 µl of cell lysate mix, with no competing plasmid, instead using increasing concentrations of the capacity monitor plasmid itself (**Supplementary Figure 1B**). This revealed that the max GFP production rate reaches a plateau at 50 nM of plasmid DNA (**Supplementary Figure 1B**). In addition, the max GFP production rate per DNA (**Figure 1C**) can be used to observe the competition for resources; a decrease in max GFP production per DNA is measured when DNA concentration is higher than 30 nM (**Figure 1C**). This decrease highlights that the amount of resources is limited for the expression of GFP from a plasmid due to competition with the pool of other copies of this plasmid.

**Figure 1:**
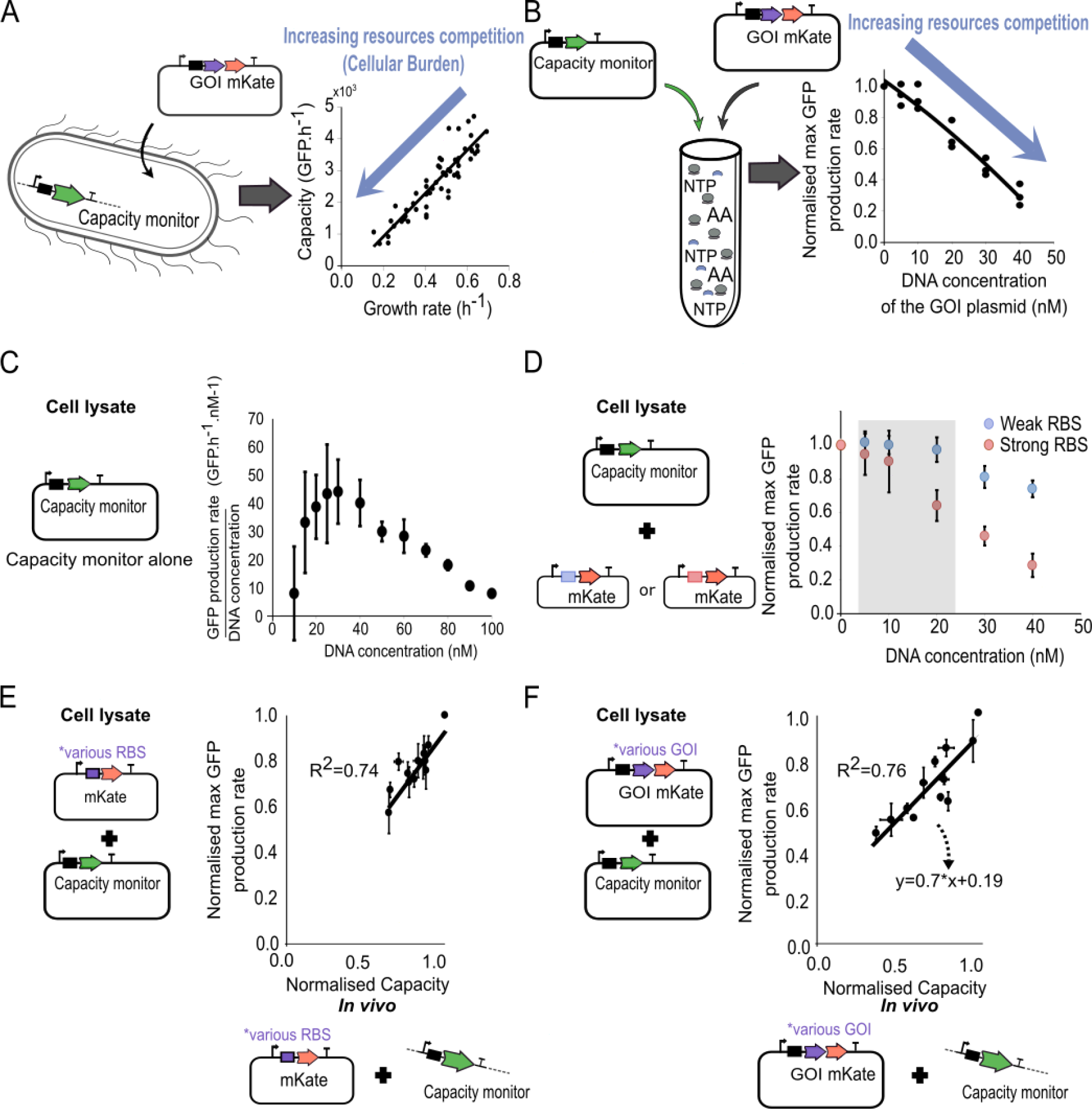
A method to measure resource competition using a capacity monitor in cell lysate. (**A**) *In vivo* resource competition in DH10B *E. coli* cells between a genome-integrated GFP capacity monitor gene and a plasmid-based gene of interest (GoI) tagged with mKate. The graph illustrates the correlation between the inferred expression capacity (measured as GFP production rate per cell) and the cell’s growth rate. (**B**) Resource competition in cell lysates expressing the capacity monitor from a plasmid and the GoI from another plasmid. The graph illustrates the correlation between the normalized max GFP production rate and the concentration of the GoI plasmid added to the lysate. (**C**) Measured max GFP production rate divided by DNA concentration with the capacity monitor plasmid added at different concentrations in a 10 µl cell lysate mix. Error bars show standard error of three independent repeats. (**D**) The normalized max GFP production rate measured in 10 µl cell lysate containing 30 nM of the capacity monitor plasmid and different concentration plasmids with *mkate* and either a strong (red) or a weak (blue) RBS sequence. The grey area represents the concentration of plasmid that leads to competition for translational resource. Error bars show standard error of three independent repeats. (**E**) Correlation between normalized max GFP production rates measured in cell lysate and normalized capacity measured *in vivo* with DH10B-GFP cells. The constructs in this experiment all express *mkate* with different strength RBS sequences. Error bars show standard error of three independent repeats. (**F**) Correlation between normalized max GFP production rate measured in cell lysate and normalized capacity measured *in vivo* with DH10B-GFP using constructs with various genes of different sizes paired with RBS BCD2. Error bars show standard error of three independent repeats.

*In vivo* in *E. coli* the main cost of gene expression is attributed to translation^6,29,30,32,33^. However, in cell lysates, while NTPs and amino acids are added in excess, polymerases, ribosomes and their associated machinery (sigma factor, tRNAs, chaperones, initiation and release factors) are added at an unknown amount and thus, the cost of transcription or translation is unknown. In order to determine the relative contributions of transcription and translation to resource competition in cell lysates, we next introduced two different plasmids to each compete with the capacity monitor plasmid. The first plasmid contains the *mkate* gene, paired with a constitutive promoter (J23106) and a strong RBS, and was used to measure the cost of both transcription and translation. The second plasmid is the same but has a very weak RBS that produces no measurable mKate protein, and so imparts a transcriptional cost and a negligible translational cost. We added different concentrations of each plasmid to the cell lysate mix along with 30 nM of the capacity monitor plasmid and measured the corresponding max GFP production rates. These values were then normalised to the max GFP production rate when the capacity monitor plasmid alone was present (*i.e*. normalised max GFP production rate = 1.0 in a cell lysate mix containing only the capacity monitor plasmid).

No decrease in max GFP production rate was observed with addition of up to 20 nM of weak RBS plasmid, implying no competition for transcriptional resources at these concentrations (**Figure 1D**). In contrast, addition of the strong RBS plasmid at these concentrations gives a significant decrease in normalised max GFP production rate (**Figure 1D**), indicating competition for translational resources in the cell lysate mix. Indeed, when 20 nM of a “competitor” plasmid is assayed with 30 nM of capacity monitor plasmid in cell lysate, translation is clearly the major cost. These conditions therefore offer a similar regime to those seen *in vivo* with *E. coli* where translation is the major cost of gene expression.

### The cost of protein production in cell lysate assays correlates with costs observed *in vivo*

We next measured the burden of a collection of plasmids expressing mKate at different levels in the cell lysate in the conditions determined above and compared these to *in vivo* measurements. We constructed a library of plasmids with *mkate* under control of the same promoter (J23106) but different RBS sequences (**Supplementary Table 1**) in order to affect burden only by altering translation initiation efficiency. These plasmids lead to a limited but measurable burden simply from overexpression of the mKate fluorescent protein and were assessed both by our previous *in vivo* capacity monitor approach (GFP production rate per cell) and by the cell lysate method (normalised max GFP production rate). Direct comparison between cell lysate and *in vivo* results yielded a linear relationship with good correlation (R^2^ = 0.74, **Figure 1E**). As the GFP production rate is a proxy of resource competition in both cell lysates and in *E. coli*, we can conclude that the competition for translational resources in the cell lysate matches those seen in *in vivo*.

Having demonstrated that cell lysate is predictive of *in vivo* burden when translation initiation rate is altered, we next looked to see if it can predict the burden of producing different proteins. First, we constructed a standard entry vector to enable the protein coding sequence of a Gene of Interest (GoI, **Supplementary Table 2**) to be rapidly cloned by Golden Gate DNA assembly into a standard format for our cell lysate assay (**Supplementary Figure 1C**). This design leads to the GoI protein coding sequence being placed under constitutive expression (J23106) and fused downstream to mKate in order to allow expression to be verified. We constructed 3 different entry vectors, each with different RBS sequences in order to give a choice of expression levels. We selected the well-characterised B0034 RBS along with 2 Bicistronic Design (BCD2 and BCD21) sequences that ensure context-free, defined levels of translation initiation ^34^.

The protein coding sequences of 7 genes with different lengths, functions and amino acid composition and 3 truncated versions of *viob* of different lengths (**Supplementary Table 2**) were all cloned into the same entry vector with the BCD2. When assayed both in cell lysate and then *in vivo* in *E. coli* a wide range of burden was observed for this collection. The capacity monitor measurements from both cell lysate and *in vivo* experiments once again showed a good linear fit, with a R^2^ = 0.76 (**Figure 1F**), demonstrating that the resource use in cell lysates of translating different proteins (and transcribing their different mRNAs) matches the resource use in *E. coli*. Similar results (linear fits) are obtained using the same measurements in cell lysate but with different growth media for the capacity characterisation *in vivo* (**Supplementary Figure 2A & B**). However, experiments done in M9 + Glucose lead to a different linear relationship between cell lysate and *in vivo* measurements (**Supplementary Figure 2B**). This can be explained by the growth-rate dependent composition of *E. coli* ^35^ (see **Supplementary Note 1, Supplementary Figure 2C & 2D**).

### A mathematical model to predict the *in vivo* burden of protein expression from cell lysate measurements

Synthetic constructs typically involve multiple heterologous proteins expressed at different rates. To predict the burden imposed by expressing genes at different rates we can use the competitive model of translation developed by Algar *et al*., previously used alongside our capacity monitor assay ^4,36^ (**Figure 2A**). This model describes the three steps of production of proteins: initial binding of ribosomes, translation elongation and ribosome release. The total number of ribosomes – the resource being competed for – is fixed in this model and is also expected to be fixed in cell lysate experiments. The binding (a_1_), unbinding (a_−1_) and initiation elongation (b_0_) rates of ribosomes on an mRNA depend on the RBS strength (See Model simulations in **Methods**). The time needed by a ribosome bound to an mRNA to fully-translate a working protein is captured by a lumped parameter γ, which represents the elongation rate. This value will vary for each GoI depending on the protein being made and how efficiently it is translated. The parameters for the capacity monitor construct in the model are the same in all our simulations (i.e. the ribosome binding rate, a_1M_ = 0.0001 rib^−1^ RBS^−1^ s^−1-1^; the ribosome unbinding rate, a_−1M_ = 200 rib-RBS^−1^ s^−1^; the translation initiation rate, b_0M_ = 1 s^−11^; mRNA amount, size_M_; and the elongation rate, γ_M_ = 1 s^−1^ see Model simulations in **Methods**, ^36^).

**Figure 2:**
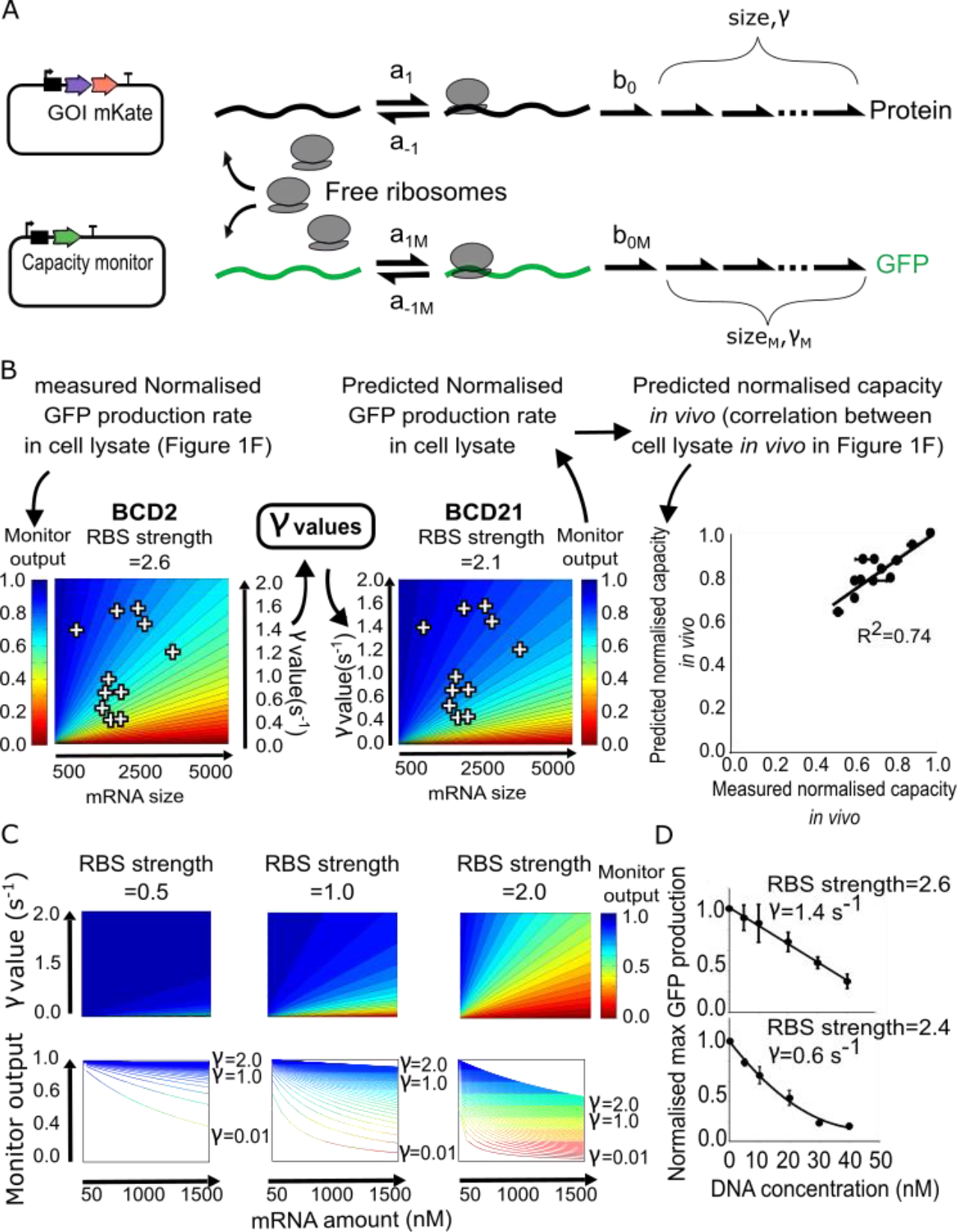
Predictive model for the competition of resources between a synthetic construct and the capacity monitor. (**A**) Overview of a mathematical model of competitive translation with a finite ribosome pool. The model represents the competition for ribosomes between the capacity monitor construct and a construct expressing a GoI. Free ribosomes bind to an unoccupied RBS at a rate a_1_ (a_1_=a_1M_*RBS strength) and either unbinds and returns to the free ribosome pool at a rate a_−1_ (a_−1_=a_−1M_/RBS strength), or initiates translation at a rate b_0_ (b_0_=b_0M_*RBS strength). Once elongation has initiated, the ribosome moves along the transcript at a rate γ. The number of elongation steps depends on “size” (mRNA size / 30 bp as it is the footprint of each ribosome on an mRNA and better represents how many can be queued on a transcript). Each elongation step is considered to proceed at the same rate γ. Once the ribosome reaches the final position on the mRNA it returns to the free ribosome pool. **(B**) Heat maps of simulated capacity monitor (monitor output) when mRNA size and the γ value of a synthetic construct are varied while RBS strength is fixed. The first heat map is used to determine the γ value of each construct used in Figure 1F. The second heat map is used to deduce the monitor output using these calculated γ values. As the prediction is done for cell lysate, the *in vivo* predictions are deduced from the correlation between cell lysate and *in vivo* measurements as per Figure 1F. Error bars show standard error of three independent repeats. (**C**) First row: heat maps of simulated capacity (monitor output) when mRNA amounts and γ values of a synthetic circuit are varied. Each heat map is a construct with a different RBS strength as denoted by the corresponding b_0_ values. Second row: capacity (monitor output) as a function of the mRNA amount. Each line corresponds to a different γ value. (**D**) Measurements in cell lysate of the normalised max GFP production rate of the capacity monitor when increasing DNA amounts of synthetic construct are added to the cell lysate assay. Top: *mkate* with a strong RBS. Bottom: *viob-mkate* with a strong RBS. The calculated γ and RBS strength values (from Supplementary Figure 3 B) are shown for each construct.

Given that it is currently impossible to determine the rate γ for a gene based on its DNA or amino acid sequence, experimental characterisation is still required to determine the burden a GoI will impart when expressed in a cell. However, our cell lysate capacity measurements now offer a rapid and standardised method to quantify this parameter without the need for cloning. To demonstrate how these data can be used to predict the burden of genes *in vivo*, we first took measurements from **Figure 1F** (expression of GoI-mKate with BCD2) and simulated them with this model. RBS strength was set to a value of 2.6 using measurements of a plasmid expressing mKate alone with BCD2 (strong translation initiation rate calculated in **Supplementary Figure 3**). Parameters in the model were set (as described in **Methods**, model simulations) in order to simulate the monitor output (equivalent of the normalised max GFP production rates in cell lysate) as a function of the mRNA size and γ value. The measured capacities and known mRNA sizes were then used to determine the γ value for each of the considered genes (**Figure 2B**). We then used this model-inferred γ value for each of these genes in an equivalent simulation but with a weaker RBS, specifically the BCD21, which we experimentally measured to correspond to RBS strength=2.1 (weaker translation initiation rate calculated in **Supplementary Figure 3**). Using this simulation, we first predicted the cell lysate normalised max GFP production rate for these new constructs, and then using the known linear relationship between lysate and *in vivo* measurements (**Figure 1F**, y = 0.7x + 0.19, with y = Normalised max GFP production and x = Normalised Capacity), we extended this to predict *in vivo* performance (**Figure 2B**). After then building and measuring the burden of this library of BCD21 constructs in *E. coli* we were able to determine that our model-based predictions of burden matched *in vivo* data with good correlation (R^2^=0.74, **Figure 2B**). Thus, with only the cell lysate data and knowledge of the mRNA length and RBS strength, we are able to predict the burden of different genes of interest expressed at different levels in *E. coli*.

Further investigation of our model shows that increasing the cost of GoI production by increasing RBS strength (a higher RBS strength implies a higher translation initiation rate and, therefore, a larger amount of ribosome binding to an mRNA) or/and decreasing γ values (a low γ value corresponds a slower global elongation rate, which leads to ribosomes staying longer on an mRNA) leads to different monitor outputs profiles as mRNA amounts are varied (**Figure 2C**). An increase of mRNA amount can be achieved through an increase of promoter strength, plasmid or gene copy number. In **Figure 2C**, the mRNA amount is used to simulate the impact of an increase of the copy number of a gene on the monitor output. At low RBS strength (e.g. 0.5), the mRNA amount and monitor output (GFP production rate) exhibit an almost linear relationship when the γ value is higher than 0.02 s^−1^ (note that the γ value for the monitor (*gfp* gene) is 1 s^−1^). In this context, the impact of the translation of several genes should be easily deduced as the burden is *additive* (*i.e*. the decrease in monitor output is the sum of the decreases in monitor output values for each gene measured individually). Even with a strong RBS (i.e. high RBS strength), a linear or close-to-linear relationship between the mRNA amount and monitor output is observed if the γ value is higher than 1 s^−1^. However, at high RBS strengths, a decrease of the γ value leads to a faster-than-linear decrease in the monitor output as mRNA amount increases (e.g. RBS strength=2 in **Figure 2C**). This means that genes that have a high cost of translation (low γ values) expressed from strong RBS sequences do not simply have additive burden, but instead will yield a nonlinear decrease in monitor output as mRNA numbers increase. Using cell lysate measurements, we experimentally demonstrated this effect by comparing the normalised max GFP production rate from the monitor construct when it competes against expression of a gene with a high γ value (*mkate*, γ = 1.4 s^−11^) and with a strong RBS (RBS strength=2.6) versus competing against expression of a gene with a low γ value (*viob-mkate*, γ = 0.6 s^−11^) and a strong RBS (RBS strength =2.4). To mimic increased mRNA levels, we simply added more DNA for these two plasmid constructs. As predicted by our model we saw a linear relationship for the high γ GoI (**Figure 2D**, upper graph), and a nonlinear relationship for the low γ GoI (**Figure 2D**, lower graph).

### Predicting the burden of *in vivo* expression of a pathway operon

As an exemplar case to predict *in vivo* burden of multiple genes, we used the four gene pathway for the biosynthesis of beta-carotene, a metabolite of interest for synthetic biology that is used industrially in nutritional supplements, cosmetics and animal feed ^37^, and offers applications in medicine too ^38^. We took the genes from *Erwinia uredovora* ^39,40^, codon-optimised these for *E. coli* and characterised the individual burden of each gene’s expression in our standard cell lysate assay using our vector with BCD2. This gave us the γ value of each enzyme (**Figure 3A, Supplementary Figure 4** and **Supplementary Note 2**). We then used MoClo Golden Gate DNA assembly ^41^ to construct a collection of 17 operons expressing all four enzymes at a variety of levels. To ensure diversity in both transcription and translation of the genes, the operons were constructed with one of three different constitutive promoters and with partially-randomised RBS sequences for each enzyme (**Figure 4A, Supplementary Figure 5A**).

**Figure 3:**
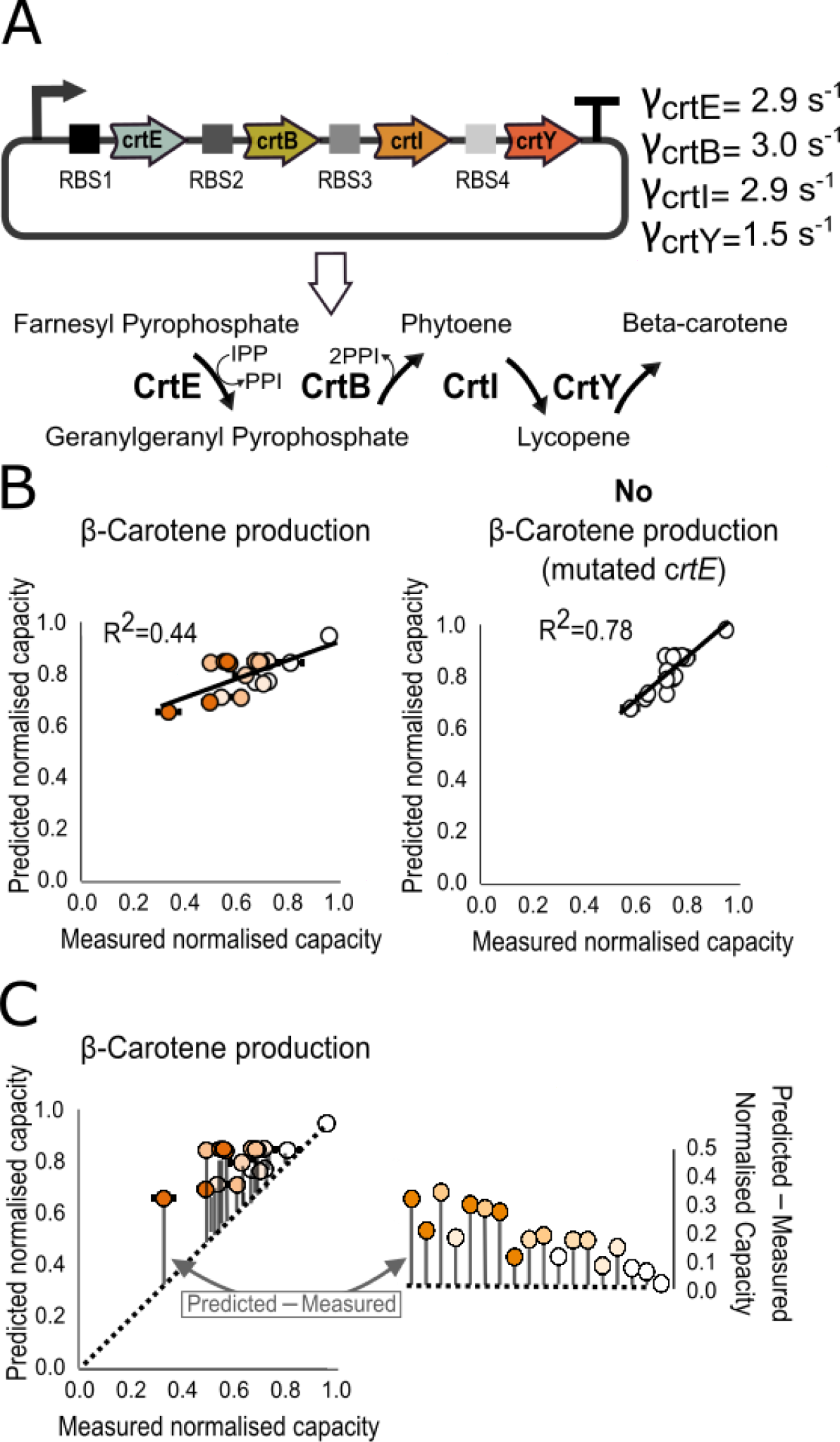
Predicting the burden of operon designs for the beta-carotene biosynthesis pathway. (**A**) Diagram of the beta-carotene pathway and the γ values for the four enzyme-encoding sequences as measured by the cell lysate capacity assay (Supplementary Figure 3). The operon is designed with partially-randomised RBS sequences and one of three different promoters: J23113 (weak), J23106 (medium) or J23100 (strong). (**B**) Model-predicted burden of each operon design compared to the measured capacity of *E. coli* expressing the operons with or without an inactivating mutation in the *crtE* gene. The orange intensity in each circle represents the beta-carotene production as level measured for each strain (see in Supplementary Figure 5). Error bars show standard error of three independent repeats (**C**) Model-predicted burden of each operon design compared to the measured capacity of *E. coli* expressing the working pathway operons (same data as panel B). The diagonal dot line represents equality between the predicted and measured normalised capacity. Grey bars indicate the difference between the predicted and measured normalised capacity of the 17 operons. The righthand plot compares the relative differences between the predicted and measured normalised capacity for the 17 operons and the strain-only control.

The model relies on 4 parameters to make predictions of the burden caused by the production of an operon: mRNA size, γ value, RBS strength and mRNA amount. The mRNA size of each enzyme is known and the γ values are determined from the cell lysate capacity assay. However, the mRNA amount and the RBS strengths are estimated using values from previous work (see **Methods**, model simulations) and using the RBS Calculator ^14^ (see **Supplementary Figure 5, Supplementary Note 3**). As the four genes in the operon exhibited a γ value higher than 1 s^−1^ (as may be expected for codon optimised genes), we assumed burden of expression would be additive and used the model to predict resource consumption for the 17 different operon versions. For the first round of predictions we made the initial assumption that the total burden of the operon *in vivo* would entirely be due to the cost of expressing the genes and that there would be no burden caused by the specific role of the genes, *i.e*. we assumed no significant burden on metabolism through conversion host metabolites into beta-carotene. However, when we compared the predicted effect on capacity from our model with subsequent measurements taken in *E. coli* (**Figure 3B**) we only saw a weak correlation (R^2^=0.44). Upon quantifying the colour of the different *E. coli* we observed a wide diversity in beta-carotene production from the different designs (**Supplementary Figure 6**), and thus it became evident that the *in vivo* burden of each operon must also relate to the metabolic cost of running the pathway in the host cell, not just the cost of expression of the enzymes. Expression of the enzymes depletes the cell of key metabolites, such as pathway precursors farnesyl pyrophosphate (FPP) and isopentenyl pyrophosphate (IPP) involved in the terpenoid backbone synthesis ^42^, and this presumably affects cell growth.

To verify this, we targeted a mutation to the active site of the first enzyme of the pathway, CrtE, in order to inactivate it and effectively cease metabolic conversion for the whole pathway (see **Methods**, construction of the beta-carotene operon). The mutation was designed to have no effect on the expression of the enzymes and so gene expression burden was still seen when the 17 operons, each with this mutation, were re-characterised *in vivo* for their effect on *E. coli* capacity. With no beta-carotene production, our predictions for the burden of each design now showed a much closer match to *in vivo* data (R^2^=0.78, **Figure 3B**). Our cell lysate-assay-based model is, thus, able to predict the impact on the host of expressing multiple genes and as expected gives the most accurate predictions when the burden is the result only of competition for gene expression resources. Interestingly, this means that our platform offers potential further use for separating “expression burden” from “metabolic burden” by simply subtracting the predicted capacity from the cell lysate data from the subsequent capacity of cells running the pathways as measured *in vivo*. The resulting difference calculated from this gives a value of burden that is not predicted to be from gene expression (**Figure 3C, Supplementary Figure 6**). How this value relates to the burden of pathway productivity, metabolite consumption and intermediate accumulation is discussed in **Supplementary Note 4**.

## Discussion

This work demonstrates that a standard cell lysate-based assay can be used to quantify the burden of expressing a protein coding sequence and provides an otherwise missing parameter for predicting the burden synthetic gene expression places on *E. coli*. Using a collection of plasmids with a range of RBS sequences and different protein coding sequences we demonstrated here that competition for translational resources in cell lysates serves as a good predictor for *in vivo* behaviour in *E. coli*. Furthermore, we provide a standard entry vector to enable quick, standardised characterisation of a GoI with cell lysates and we accompany this with a mathematical model that enables prediction of the *in vivo* performance from cell lysate measurement data. The cell lysate assay effectively quantifies a global parameter, we here term γ, which is the combination of all sequence-dependent parameters (nucleotide composition, secondary structure of the mRNA, translational pausing, presence of rare codons or use of rare amino acids) involved in the resource costs for translating a protein coding sequence. Quantifying this is necessary for accurate approximation of the burden of protein expression as there is currently no *in silico* tool to predict the cost, time or efficiency of translation for a given RNA or amino acid sequence. When combined with estimates of RBS strength from the RBS Calculator ^14^, these cell lysate measurements enable predictions both for single genes when there are changes in the translation efficiency, and for multigene systems such as the operon example here.

Another exciting finding from this work is the possibility that our platform can separate “expression burden” from “metabolic burden”, something that cannot easily be done *in vivo* due to the combined effects that all types of burden have on host cell growth rate. Our characterisation of beta-carotene pathway operons demonstrates that these two types of burden are jointly-responsible for decreased growth rates of hosts expressing heterologous genes to produce metabolites. Most methods for metabolic pathway optimisation seek to produce the most product whilst doing so with the minimal cost of expression of the enzymes ^43,44,45^. Quantifying the individual contributions to burden of both gene expression and pathway productivity offers a new tool for designing the most productive pathways and investigating the mechanistic causes of burden in more depth.

To further deconstruct the mechanisms of expression burden and quantify resource use at a molecular level, future iterations of our cell lysate approach could make use of defined *in vitro* expression systems such as PURE Express (NEB), which contains known quantities of purified components such as polymerases and ribosomes ^46^. Full control of the make-up of cell lysates would provide a route towards determining the main components that are required for efficient gene expression and could be used to investigate which factors are limiting for different genes. For example, charged tRNAs may be limiting for genes with rare codons, while chaperones may be limiting for genes requiring complex folding. Such an approach would likely reveal hidden mechanisms and constraints in gene expression, highlighting basic components to increase in cells when needing to efficiently overexpress certain genes, while also providing a more complete list of the components needed for the construction of minimal cells.

Further exploration with cell lysates and/or the PURE system will likely reveal the key molecular interactions involved in the translation elongation and release processes, and ideally enable the development of a biophysical model suitable for predicting the protein cost based only on the input of a DNA sequence. Such a predictor will complete the set of tools (with transcription [21] and translation initiation [22] predictors) necessary to design and develop reliable circuits *in silico* and accelerate the construction of large and more complex systems and pathways using synthetic biology.

## Methods

### Strains and Growth media

Plasmids were transformed using standard procedures ^47^ in chemically competent *E. coli* DH10B-GFP. DH10B-GFP is the DH10B strain with genome integrated capacity monitor that consists of a strong constitutive promoter (J23100), strong synthetic RBS (tactagagaaatcaaattaaggaggtaagata), a codon-optimized superfolder GFP ^48^ coding sequence, and a synthetic unnatural bidirectional terminator integrated in the λ loci of *E. coli* genomes Bacterial growth was performed at 37°C in minimal media M9 supplemented with 0.5% fructose (or 0.5% glucose or pyruvate, see **Supplementary Figure 2**) and chloramphenicol (35 μg/ml).

### Construction of the GoI-mkate library

The high-copy plasmid pSB1C3 (BioBricks Foundation, Supplementary Data), chloramphenicol-selectable, was used as a backbone to construct the standard entry vector for GoI insertion (**Supplementary Figure 1C**). To construct the platform: we first PCR amplified pSB1C3 (forward primer: taagccagccccgacacccg / reverse primer: tgaaccacagagtgattaat) and *lacZ* under control of Plac promoter flanked by BsaI restriction sites (forward primer: gcagctggcacgacaggttt / reverse primer: ttatgcggcatcagagcaga). Second, the linker-mkate sequence was codon optimised and ordered on GeneArt with the RBS sequences (BCD2 and B0034). The different parts were then assembled and cloned using the Gibson Assembly method ^49^ to obtain the standard entry vector described in **Supplementary Figure 1C**.

The GoI (**Supplementary Table 2**) were all obtained using biobricks of the iGEM Parts Registry as template and PCR amplified to be flanked by the proper BsaI restriction sites (ggtctcannnn). Golden Gate assemblies were set up by pipetting 40 fmol of backbone and insert, 0.5 µl of BsaI (NEB UK), 0.5 µl of T7 DNA ligase (NEB UK), 1 µl T4 buffer (NEB UK) and completed with water for final volume of 10 µl. Then the mix was put in a thermocycler for 30 following cycles: 42°C for 2 minutes / 6°C for 5 minutes / 55°C for 1 hour / 80°C for 10 minutes.

### Construction of the beta-carotene operons

The beta-carotene operons were build using MoClo toolkit ^41^. The Level 0 library is composed of constitutive promoters of the Anderson collection (J23114, J23113, J23100, J23106 and J23115), a random collection of RBSs, the 4 enzymes of the beta-carotene pathway and the terminator T1 ^41^. The level 1 was designed to put each enzyme under the control of a random RBS and to place the genes of the beta-carotene operons in the following order at the final level: *crtE*, *crtB*, *crtI* and *crtY*. The cloning was done using Golden Gate assembly as previously described, and then transformed in DH10B-GFP.

Two constructs (B3-1 str and B10-2 str, containing promoter J23100) were obtained by PCR amplification to introduce mutations in the “weak” promoter sequence J23113 (phosphorylated forward primer: tacggctagctcagtcctaggtatagtgctagcgcaagggcccaag reverse primer: ttcacagagtggcctcgtga) of previously obtained constructs (B3-1 and B10-2). The resulting PCR fragments were ligated using T4 DNA ligase and transform in DH10B-GFP.

All the mutated *crtE* constructs were obtained by PCR amplification (forward primer: gccgctatgccctgcatggacg, reverse primer: cgcggcttcgctgatcctt) and T4 ligation to introduce mutations in *crtE* leading to an inactivated CrtE enzyme. The mutation of *crtE* (from gacgat to gccgct) was chosen in order to modified the active site of *crtE* deduced by sequence homology of *crtE* sequences from *Erwinia uredovora* ^50^, *Erwinia herbicola* ^50^, *Rhodobacter capsulatus* ^50^, *Arabidopsis thaliana* ^52^ and *Euglena gracilis* ^52^.

### Cell lysate mix preparation and reactions

The cell lysate preparation is based on the protocol of Sun *et al*. ^28^. The protocol was modified by using sonication ^53^ instead of beads beater to obtain DH5 alpha cell extract. After washing the cells as described in Sun *et al*. protocol (Day 3 step 18 in ^28^) with S30A buffer (14 mM Mg-glutamate, 60 mM K-glutamate, 50 mM Tris 2mM DTT, pH 7.7), the cells were centrifuged 2000g for 8 min at 4°C. The pellet was re-suspended in S30A (pellet mass (g)*0.9 ml). The solution was split in 1ml aliquots in 1.5 ml Eppendorf tubes. Eppendorf tubes were placed in a cold block and sonicated using a Vibra-Cell™ Ultrasonic Liquid Processors VCX 130 using the followings procedure: 40 s ON – 1min OFF – 40 s On −1min OFF – 40 s ON. Output frequency 20 kHz, amplitude 50%.

The remaining protocol follows the procedure described in ^28^ for Day 3, Step 37. mRNA and protein synthesis are performed by the molecular machineries present in the extract, with no addition of external enzymes. The Amino Acid Solution and Energy Solution described in ^28^ was added to the cell extract. Thus, 3 Mg-glutamate, 85 mM K-glutamate, 1.5 mM each amino acid except leucine, 1.25 mM leucine, 50 mM HEPES, 1.5 mM ATP and GTP, 0.9 mM CTP and UTP, 0.2 mg/mL tRNA, 0.26 mM CoA, 0.33 mM NAD, 0.75 mM cAMP, 0.068 mM folinic acid, 1 mM spermidine, 30 mM 3-PGA, 2% PEG-8000 were added to the extract to obtain the final cell lysate. Reactions took place in 10 μL volumes at 29°C in 384-well plate.

### Burden assay *in vivo* and data analysis

For burden measurements in DH10B-GFP, cells were grown at 37 °C overnight with aeration in a shaking incubator in 5 ml of defined supplemented fructose M9 media with chloramphenicol (35 μg/ml). In the morning, 20 μl of each sample was diluted into 1 ml of fresh medium and grown at 37°C with shaking for another hour. We then transferred 200 μL OD_600_ into a 96-well plate (Costar), placed samples in a Synergy HT Microplate Reader (BioTek) and incubated them at 37°C with orbital shaking at medium setting, performing measurements of GFP (excitation (ex.), 485 nm; emission (em.), 528 nm), RFP (ex., 590 nm; em., 645 nm), OD (600 nm) and OD (700 nm) every 10 min

Growth were calculated using OD_700_ with: growth rate at t2= [ln(OD(t3)) – ln(OD(t1))] / (t3 – t1), with t2 = time of the mid exponential phase, t3 = t2 + 0.5 hr and t1 = t2 – 0.5 hr.

Protein production rates per hour were calculated with:

GFP production rate at t2 = [(total GFP(t3) – total GFP(t1)) / (t3 – t1)] / OD(t2), and RFP production rate at t2 = [(total RFP(t3) – total RFP(t1)) / (t3 – t1)] / OD(t2).

The normalised capacity stands for the GFP production rate measured in strains with the DH10B-GFP containing a plasmid divided by the GFP production rate measured in DH10B-GFP without any plasmid.

### Resource competition assay in cell lysate and data analysis

For resources competition in cell lysate, reactions took place in 10 μL volumes at 29°C in 384-well plate (Nunc™ 384-Well). Each reaction is a mix of 7.88 μL cell lysate (cell extract + amino acid + energy solution + Mg-glutamate buffer + K-glutamate buffer + PEG, see ^28^), plasmid DNA and complete with sterile water to get 10 μL. Plates were centrifuged 5 min, 2000 rpm at 4°C (Eppendorf, centrifuge 5810R). Samples were placed in a Synergy HT Microplate Reader (BioTek) and incubated them at 37°C with orbital shaking at low setting, performing measurements of GFP (excitation (ex.), 485 nm; emission (em.), 528 nm), RFP (ex., 590 nm; em., 645 nm) every 5 min.

Protein expression rates per hour were calculated with:

GFP production rate at t2 = (total GFP(t3) – total GFP(t1)) / (t3 – t1), and RFP production rate at t2 = (total RFP(t3) – total RFP(t1)) / (t3 – t1).

The maximal expression rate value was selected as described in **Figure 1C**. The normalised max GFP production rate stands for the max GFP production rate measured in a cell lysate mix containing the capacity monitor plasmid and another plasmid divided by the max GFP production rate measured in a cell lysate mix with only the capacity monitor plasmid.

### Beta-carotene measurements

*E. coli* were incubated with aeration in a shaking incubator in 5ml of Minimum media M9 supplemented with 0.5% fructose at 37°C during 24 hours. Cells were harvested using centrifugation at 4000 rpm for 10 minutes (Eppendorf, centrifuge 5810R). Pellet was re-suspended in 300 μL acetone, homogenised by vortexing and incubated at 55°C for 15 min. Supernatant was collected after 1 min centrifugation at 14000 rpm (Eppendorf, centrifuge 5424). 100 μL of water was added to 100 μL of samples and OD (450 nm) was measured in a Synergy HT Microplate Reader (BioTek).

### Model simulations

We used the competitive model of translation developed by Algar *et al*. ^36^. Simulations were done using the key parameters obtained from Bionumbers.org and calculated in ^35^ along with the relative strengths of the RBSs used as determined from our experimental characterization. As J23100, J23106 and J23113 are known to be strong, medium and weak promoters, we assumed each copy of the promoter would produce about 30, 10 and 3 copies of mRNA per DNA molecules respectively (estimated from Bionumbers I.D. 107667). We added 20 nM of each plasmid in our mix leading to 600, 200 and 40 nM of mRNA in the cell lysate mix. Parameters for simulation of construct expression and capacity monitor expression were as follows: total available ribosomes, 2,500 nM (25000 ribosomes per cell ^35^ extracted from 1.6.10^12^ cells, stocked in a final cell extract volume of 9600 µl and diluted 3 time in the final cell lysate mix); capacity monitor length, 24 (720 bp /30); capacity monitor mRNAs, 900 nM (30 nM * 30 mRNA per DNA as J23100 is a strong promoter); capacity monitor b_0M_=1 s^−1^; capacity monitor γ=1 s^−1^. The RBS strength of each construct is a relative value compare to the monitor RBS strength (RBS strength of monitor = 1). For each construct the binding (a_1_), unbinding (a_−1_) and elongation initiation (b_0_) rates are function of the RBS strength and of a_1M_, a_−1M_ and b_0M_ respectively. More specifically, in our model we define a_1_ = a_1M_*RBS strength, a_−1_ = a_−1M_/RBS strength, and b_0_ = b_0M_*RBS strength ^36^.

For the sake of clarity b_0_ and γ units are noted in s^−1^ in the manuscript but are more precisely in (10 codons).s^−1^. The b_0_ and γ units are (10 codons).s^−1^ because 10 codons is approximately the footprint of each ribosome on an mRNA and thus for our model it better represents how many ribosomes can be queued on a transcript.

## Acknowledgements

OB was supported by EPSRC grant EP/M002306/1. GBS was supported by EPSRC Fellowship EP/M002187/1 and grant EP/P009352/1. TE was supported by EPSRC Fellowship EP/M002306/1 and grant EP/J021849/1. We thank Brooke Rothschild-Mancinelli for assistance with pellet photography and Francesca Ceroni for critical reading of the manuscript.

---

**Supplementary Figure 1:**
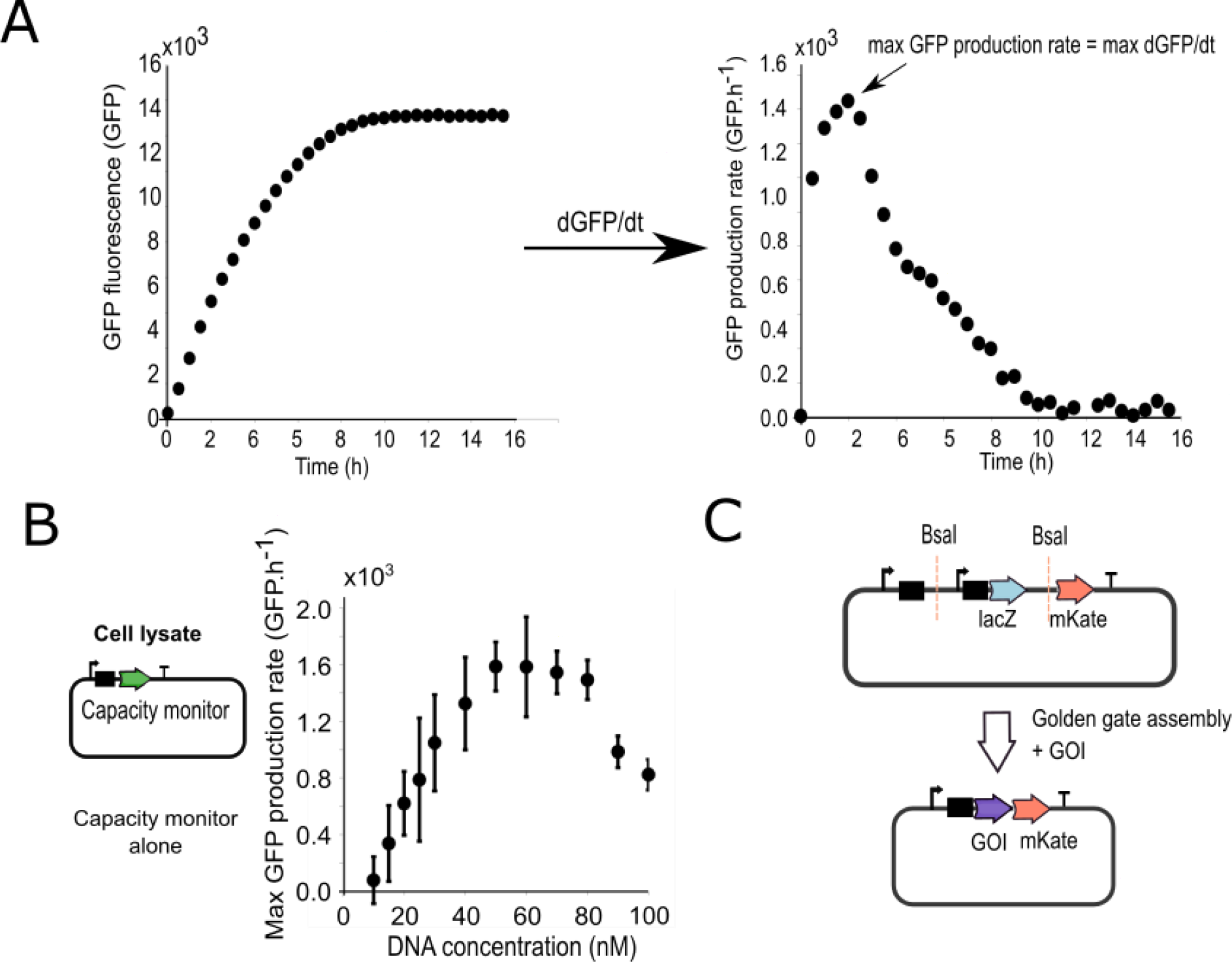
Cell lysate measurements and backbone design (**A**) The graph on the left is an example of the GFP fluorescence measurements in cell lysate as a function of time using 40 nM of the capacity monitor plasmid cell lysate mix. The graph on the right is the derivate of the GFP fluorescence, i.e. the GFP production rate. GFP production rates were calculated with: GFP production rate at t2 = [total GFP(t3) – total GFP(t1)]/(t3 – t1) with t2 = time of the measurement, t3 = t2 + 0.25 hr and t1= t1 – 0.25 hr. The maximum GFP production rate is the maximum value of GFP production rate. (**B**) Measured max GFP production rate with the capacity monitor plasmid added at different concentrations in a 10 µl cell lysate mix. Error bars show standard error of three independent repeats. (**C**) The entry vector design for this work consists of a promoter J23106, the BCD 2 or BCD 21 or RBS B0034, a *bsa*I restriction site (ggtctcccatt), promoter pLac, lacZ, a *bsa*I restriction site, linker sequence (aggtctcagctt), codon optimised mkate and terminator. Golden Gate assembly was used to integrate different GoIs leading to GoI-mKate fusion proteins.

**Supplementary Figure 2:**
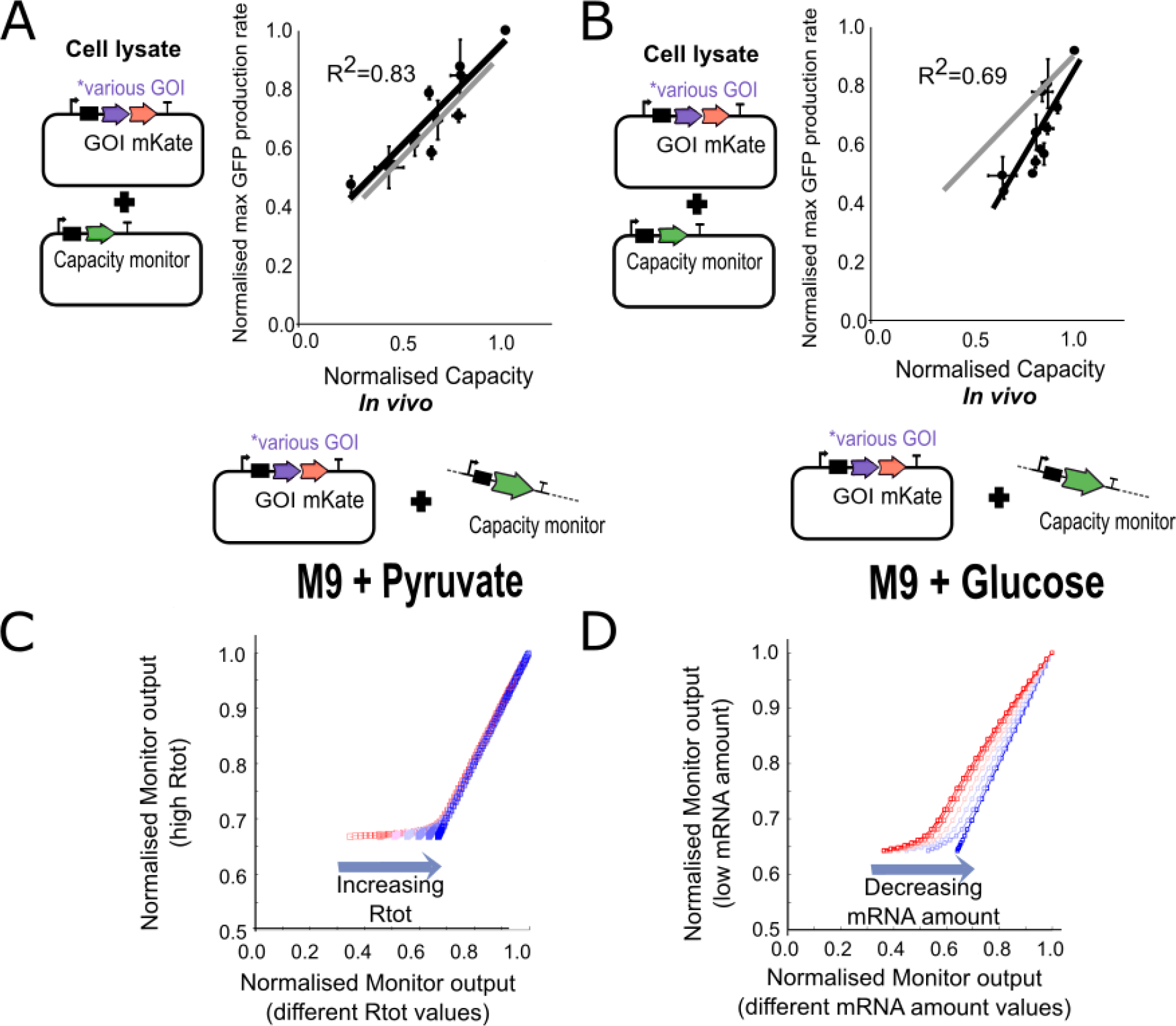
Medium-dependent burden. (**A**) Correlation between normalized max GFP production rates measured in cell lysate (same value as Figure 1E) and normalized capacity measured *in vivo* with DH10B-GFP using constructs with various genes of different sizes paired with RBS BCD2. All *in vivo* measurements have been done with the same strain as Figure 1E but grown in minimal media + 0.5% pyruvate. Error bars show standard error of three independent repeats (**B**) Correlation between normalized max GFP production rate measured in cell lysate (same value as Figure 1E) and normalized capacity measured *in vivo* with DH10B-GFP using constructs with various genes of different sizes paired with RBS BCD2. All *in vivo* measurements are with the same strain as Figure 1E but grown in minimal media + 0.5% glucose. Error bars show standard error of three independent repeats. (**C**) Simulations of the Monitor output in competition with circuits with different γ values with Ribosome fixed at 50000 (y axis). Simulations of the Monitor output in competition with circuits with different γ values with Ribosome fixed at 50000, 10000, 5000, 2000 or 1000 nM (x axis) (**D**) Simulations of the Monitor output in competition with constructs with different γ values with mRNA fixed at 100 (y axis). Simulations of the Monitor output in competition with constructs with different γ values with mRNA fixed at 5000, 4000, 3000, 2000, 1000, 500 or 100 nM (x axis).

**Supplementary Figure 3:**
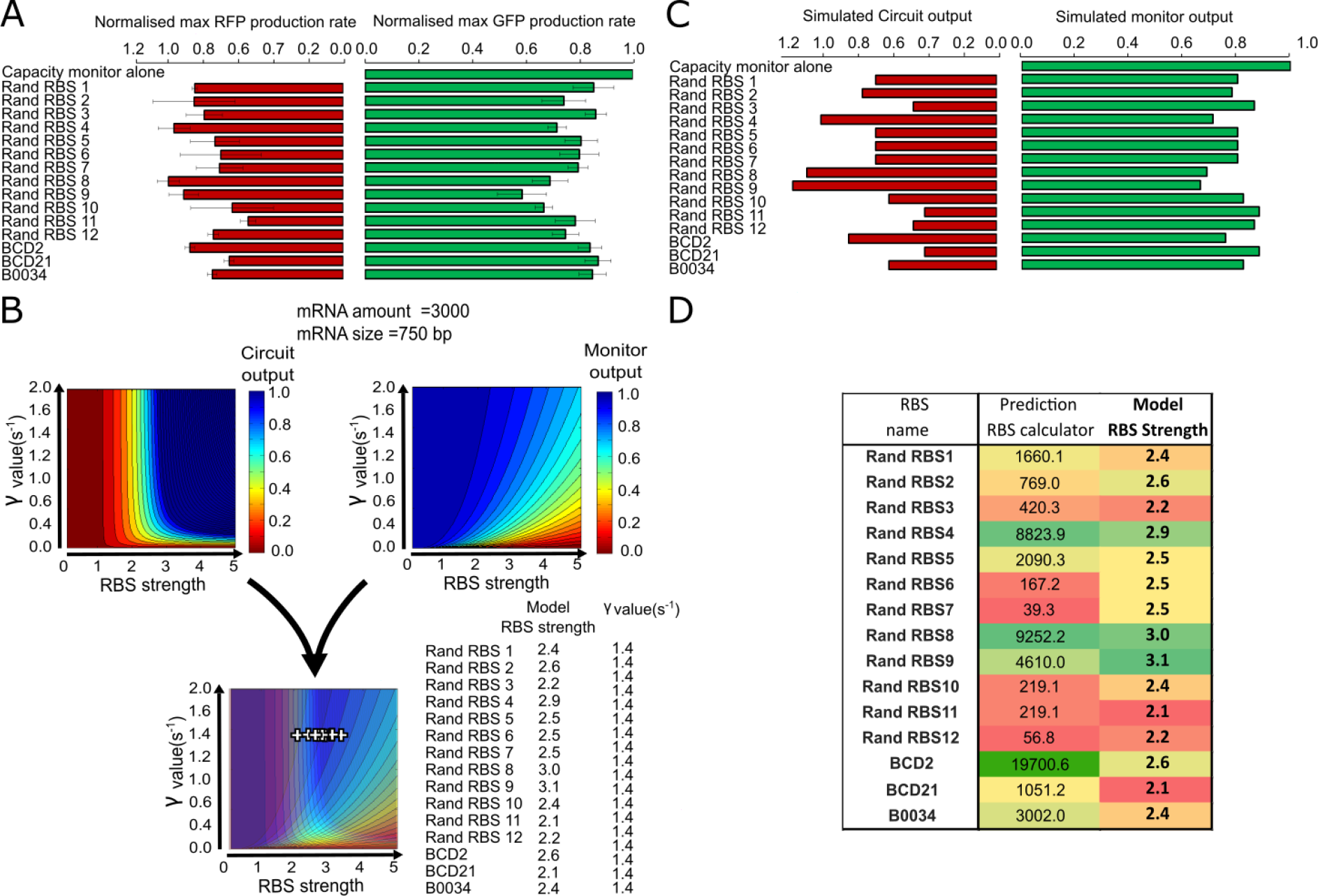
Cell lysate measurements and simulated output and capacity at steady state determined by a translational resource model. (**A**) Normalised max RFP production rate and normalised max GFP production rate measurements of constructs with mkate alone under control of different RBS/BCDs (library from Figure 1E). Normalised max RFP production rates are the max RFP production divided by the mean max RFP production of the Rand RBS 8 construct (strongest of the collection). Normalised max GFP production rates are the max GFP production divided by the max GFP production of capacity monitor expressed alone in the cell mix as described in Methods. Error bars show standard error of three independent repeats. (**B**) Simulation of the construct output (corresponding to the normalised max RFP production rate measurement) and monitor output (corresponding to the Normalised max GFP production rate measurement). The construct and monitor outputs graphs have been merged to deduce RBS strengths and γ value of the mkate construct library. All the constructs produce the same mkate protein and thus have the same γ value. (**C**) Results of the simulation of panel B. (**D**) Comparison between the RBS calculator predictions and our model simulations. The order of RBSs strength is respected when compared between RBS Calculator predictions and our model simulations. Our simulations reveal that the strongest RBS of our collection has a RBS strength=3.0, medium RBSs have a RBS strength=2.5 and the weakest RBS has a RBS strength=2.0.

**Supplementary Figure 4:**
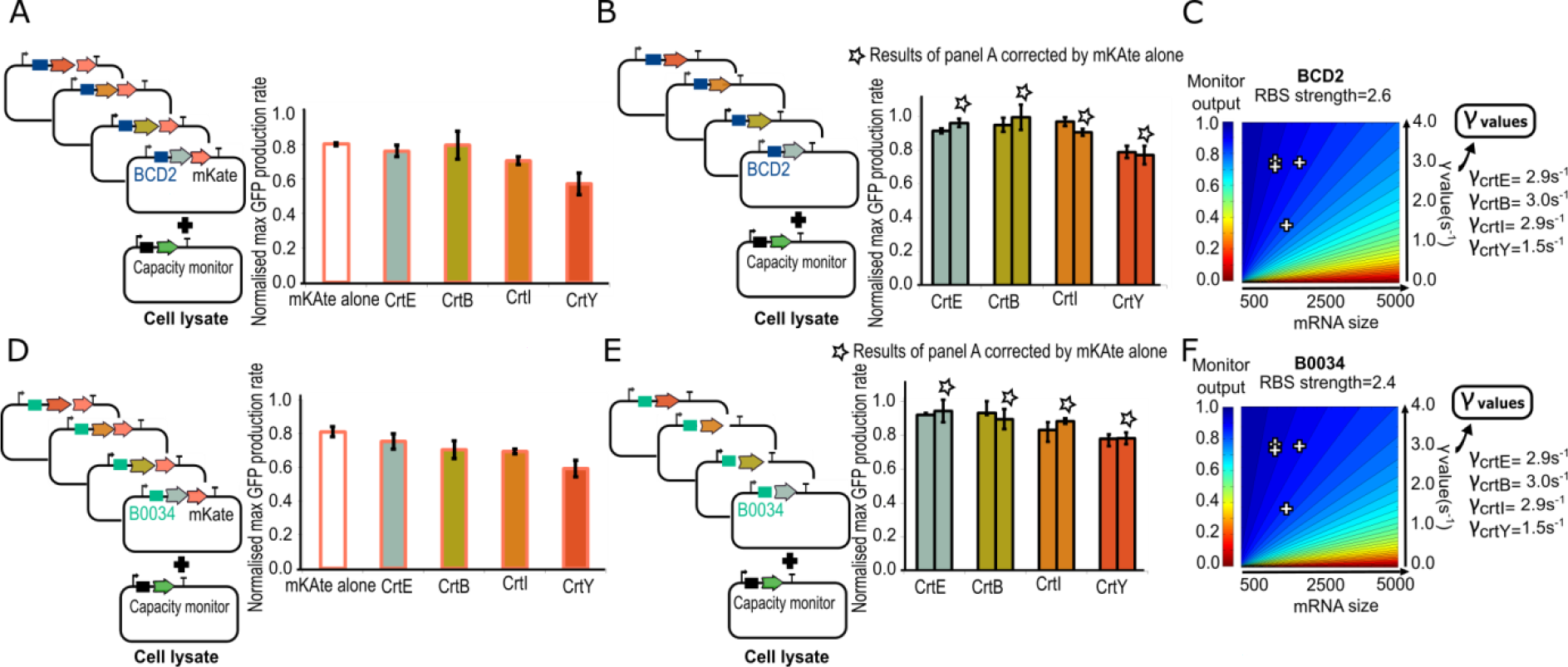
Competition for the resources in cell lysate between the enzymes of the beta-carotene pathway and the capacity monitor. (**A**) Normalised max GFP production rate measured with capacity monitor and the crt-mkate fusion added in the cell lysate mix. Crt are the enzymes of the beta-carotene pathway. The production of mKate (mKate alone) under control of BCD2 leads to a competition for the resources as the normalised max GFP production rate is lower than 1. (**B**) Normalised max GFP production rate measured with capacity monitor and the crt enzymes in the cell lysate mix. The mKate CDS has been removed to obtain the competition for resource due to the crt enzyme alone. The normalised max GFP production rate of the crt-mkate fusion corrected by removing the cost of mkate calculating using the data from panel A. The normalised max GFP production rates of the crt enzymes and of the corrected crt-mkate exhibit similar values showing that both backbone with mkate fusion and without mkate can be used for γ value calculation. (**C**) Heat maps of simulated capacity (monitor output) when mRNA size and the γ value of a synthetic construct are varied as RBS strength is fixed (the BCD2 RBS strength has been obtained as explained in Supplementary Figure 2B). (**D, E, F**) Similar experiment as panels A, B, C but with RBS B0034.

**Supplementary Figure 5:**
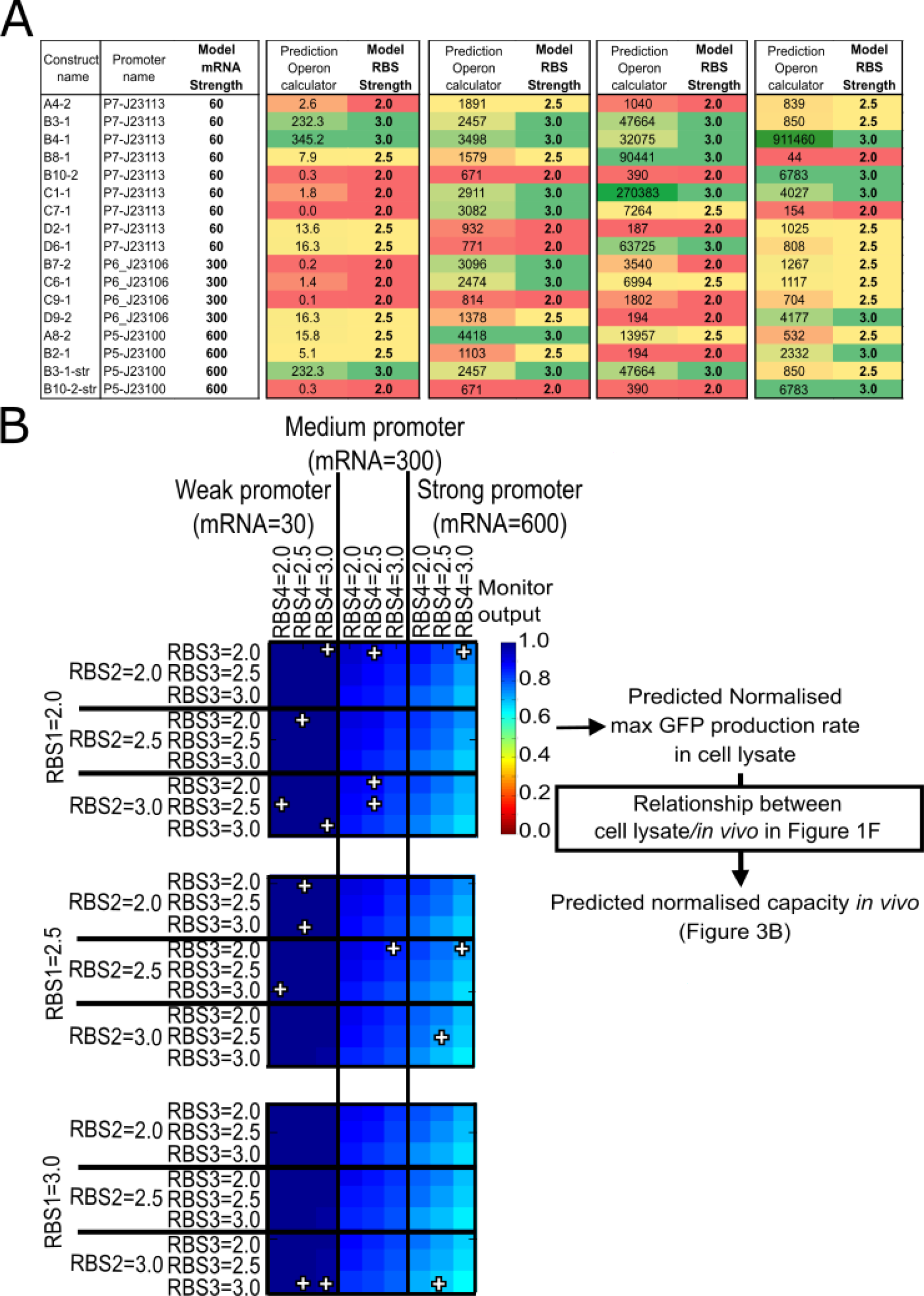
Burden prediction for beta-carotene operons. (**A**) RBS Calculator prediction using the operon calculator. We ranked the model RBS value as described in Supplementary Note 2. (**B**) Predictions from the competition between the capacity monitor and the 4 *crt* enzymes using the γ values obtain in Supplementary Figure 3 and the RBS values obtain in panel A.

**Supplementary Figure 6:**
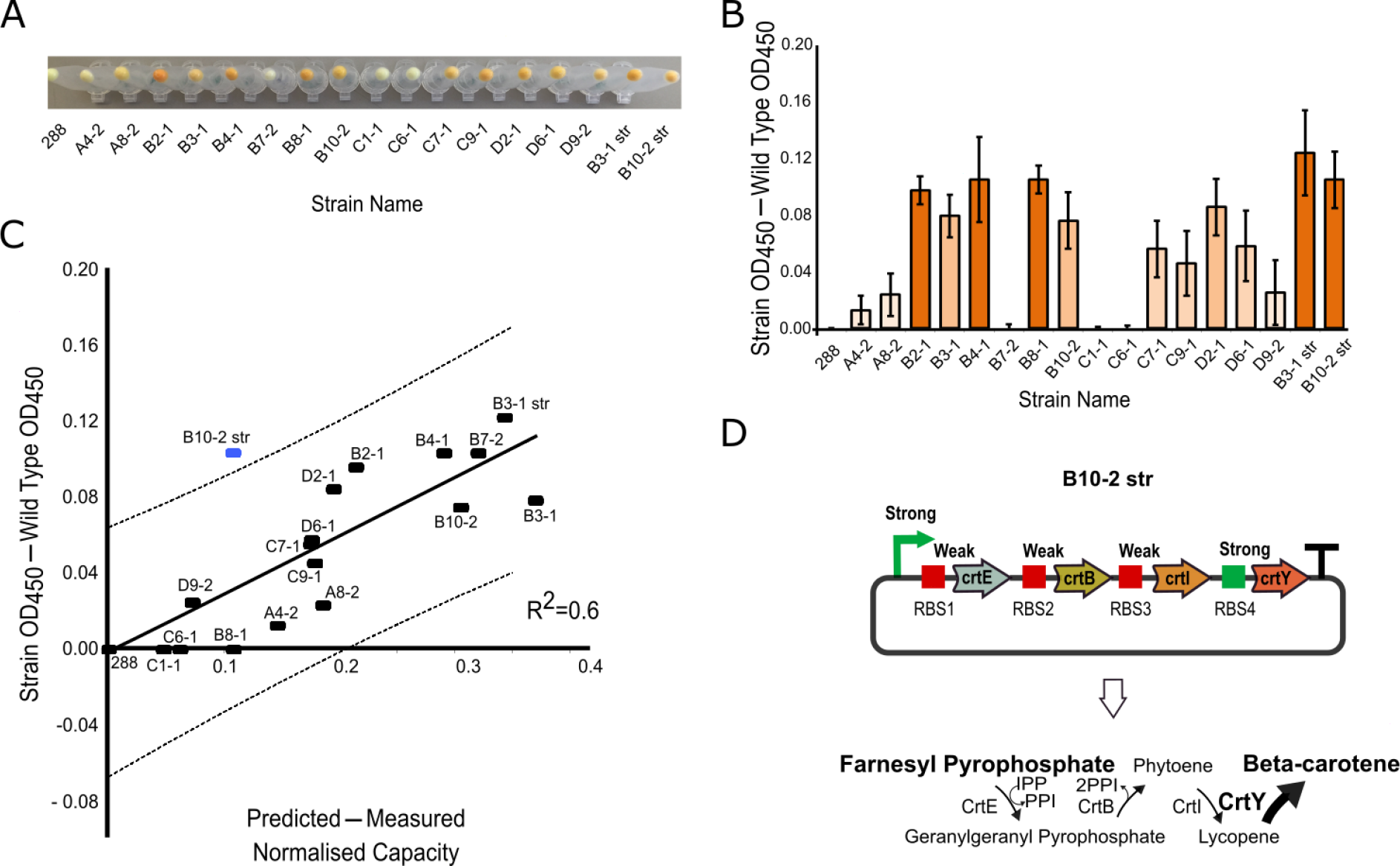
Beta-carotene production.(**A**) Pellet of the strains producing beta-carotene with the constructs of the beta-carotene operon library (**B**) Beta-carotene values obtain at OD_450_ measurements after acetone treatment (**C**) Comparison between the beta-carotene production (panel B) and the difference between the predicted and measured capacity (Figure 3C). The dotted lines represent the 95% confident interval. B10-2 str appears to be an outlier (blue dash). (**D**) Diagram of the beta-carotene pathway of the construct B10-2 str. B10-2 str is a construct with a strong promoter, weak RBS1/RBS2/RBS3 and a strong RBS4 (Supplementary Figure 5A). This construct should produce a higher amount of CrtY, the last enzyme of the beta-carotene operon, than CrtE, CrtE and CrtI.

**Supplementary Table 1:**
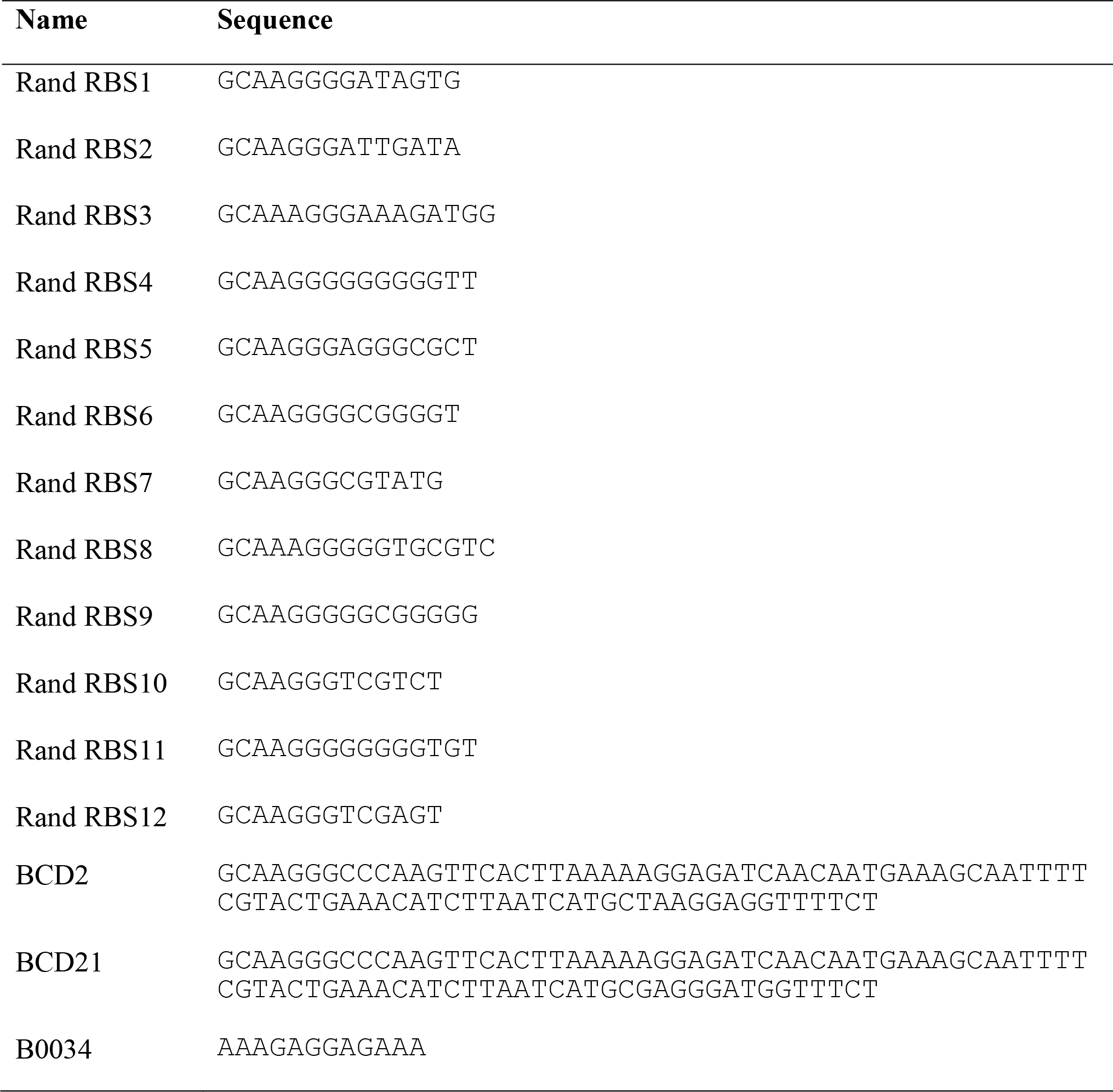
Library of BCDs/RBSs.

**Supplementary Table 2:** Library of selected Genes of Interest (GoI)

| **Name** | **Function** | **Size**<br>**(bp)** |
| --- | --- | --- |
| *empty* |  | 0 |
| *Viob_truncated3* |  | 500 |
| *suhB* | Inositol-1-monophosphatase (use for the production of glucaric Acid) | 804 |
| *URA3* | Orotidine 5'-phosphate decarboxylase (ODCase), catalyzes the synthesis of pyrimidine ribonucleotides | 820 |
| *Viob_truncated2* |  | 1000 |
| *Leu2* | 3-isopropylmalate dehydrogenase | 1111 |
| *aspA* | aspartate ammonia-lyase | 1479 |
| *atf1* | Alcohol acetyltransferase | 1572 |
| *Viob_truncated1* |  | 2000 |
| *ste12* | Pheromone-responsive transcriptional activator. | 2070 |
| *Viob* | involved in dTDP-N-acetylviosamine synthesis | 2994 |

## Supplementary Note 1: Medium-dependent burden (Supplementary Figure 2)

As the translation and transcription machinery concentrations are condition-dependent *in vivo*, we anticipate that the competition for resources will vary when experiments are done in different conditions ^6355455^. We can see some evidence of this in Figures 1E and 1F as the slopes of the linear fits indicate that zero capacity *in vivo* would still lead to some non-zero GFP expression in cell lysate. Interestingly, when we take the library of constructs expressed in DH10B in Figure 1F and measure their capacity in different growth media, we still observe a linear relationship between cell lysate and *in vivo* measurements. However, the slopes of these linear relations are different (Supplementary Figure 2). This linear relation is the same in M9 + pyruvate as in M9 + fructose, but significantly different when M9 + glucose is used. This difference is likely explained by the dependence of cell physiology on growth rate ^6355455^. Indeed, wild-type cells in M9 + pyruvate and M9 + fructose have similar growth rates (respectively 0.54 +/-0.05 h^−1^ and 0.62 +/-0.04 h^−1^). However, when cells are grown in M9 + glucose, they exhibit a considerably faster growth rate (0.85 +/-0.05 h^−1^). Using our computational model, we simulated the impact of different GoI with different γ value on the monitor output. First, we compared a condition with a high concentration of ribosomes (50000 nM, y axis on Supplementary Figure 2C) to several conditions with lower concentration of ribosomes (10000 nM, 5000 nM, 2000 nM and 1000 nM, x axis on Supplementary Figure 2C). Second, we compared a condition with a low concentration of mRNA (100 nM, y axis on Supplementary Figure 2D) to several conditions with higher concentration of mRNA (5000 nM, 4000 nM, 3000 nM, 2000 nM, 1000 nM, 500nM and 100 nM, x axis on Supplementary Figure 2D). Based on these model simulations, we predict that a decrease in mRNA amount yields a change of the monitor profile that is similar to that experimentally observed in Supplementary Figure 2B. This prediction requires further experimental results to be validated.

## Supplementary Note 2: Competition for resources in cell lysate between the enzymes of the beta-carotene pathway and the capacity monitor (Supplementary Figure 4)

Here two backbones have been used to integrate selected GoIs. These backbones contain BCD2 or BCD21, which control the expression of their corresponding GoI after Golden Gate assembly. The BCD (Bicistronic Design) is a 84 nucleotides sequence containing a first Shine-Dalgarno sequence followed by a stop codon to disrupt mRNA structures and a second Shine-Dalgarno sequence to initiate translation of the GoI. BCDs have been designed to be independent of the GoI sequence ^56^. During this work, we noticed some difficulties during cloning with BCD-containing plasmids. Indeed, half of the colonies obtained after Golden Gate assembly contained a backbone re-ligated on itself without any GoI. Thus, we decided to also provide to future users a backbone with the simple RBS B0034, as B0034 has been demonstrated previously to be largely context-independent ^56^. Measurements show similar results with B0034 but without the cloning difficulties observed with BCDs (Supplementary Figure 4).

## Supplementary Note 3: Burden prediction for beta-carotene operons (Supplementary Figure 5)

We observed in Supplementary Figure 3D that the predictions of the RBS Calculator and the model-calculated RBS strengths exhibit similar ranking order. In addition, the strongest RBS of our collection has a RBS strength=3.0, medium RBSs have a RBS strength=2.5 and the weakest RBS has a RBS strength=2.0 (obtained from the simulations shown in Supplementary Figure 3B). We used this ranking approach and the RBS strength values to estimate the model RBS strength of the sequenced RBSs from our beta-carotene operon library (Supplementary Figure 5A). First, we used the Operon Calculator of the Salis Lab (https://salislab.net/software/) to associate a value to each RBS ^14^. Second, we ranked each RBS based on the Operon Calculator predictions and on their positions within the operons. Third, we associated a model RBS strength of 2.0, 2.5 or 3.0 depending on the Operon Calculator RBS strength rank (Supplementary Figure 5A). This choice of RBS strengths appears to be accurate as the predictions based on our model-based simulations lead to a good correlation with experimental results (R^2^= 0.78, Figure 3B).

## Supplementary Note 4: Beta-carotene production (Supplementary Figure 6)

In order to quantify the metabolic burden caused by the production of beta-carotene, we assumed that the decrease of capacity was the result of the addition of the expression burden and metabolic burden. As the predicted burden using cell lysate and our model (Figure 3B) estimate the expression burden, we determined the metabolic burden from the difference between the predicted capacity and the measured capacity in *E. coli* strains expressing the original operons (i.e. without the inactivating mutation in the *crtE* gene, Figure 3C). This difference was compared to the beta-carotene production in the different strains (Strain OD_450_-wild type OD_450_ in Supplementary Figure 6B). Most of the strains fit on a linear relationship where more beta-carotene production corresponds to an apparent increase in metabolic burden, demonstrating the relevance of our method (Supplementary Figure 6C). However, as the anticipated metabolic burden should come not only from the production of beta-carotene (the pathway end product) but also from production of the intermediates geranylgeranyl pyrophosphate, phytoene and lycopene, we expect that using only the beta-carotene production as a proxy underestimates the metabolic burden in most cases. Interestingly, construct B10-2 str (blue dash in Supplementary Figure 6D) stands out as the strain expressing this produces more beta-carotene product than the small calculated metabolic burden suggests. B10-2 str is a construct with a predicted strong promoter, weak RBS1/RBS2/RBS3 and strong RBS4 (Supplementary Figure 5A), which should lead to high expression of CrtY, the last enzyme of the pathway, and low expression of CrtE, CrtB and CrtI (Supplementary Figure 6D). B10-2 str should therefore accumulate a low amount of intermediates and convert metabolites into beta-carotene more efficiently (*i.e*. with less metabolic burden). So as this appears as an outlier in the data it reaffirms that the other constructs are likely to be generating unmeasured metabolic burden via intermediate accumulation. Future work could quantify the pathway intermediates in constructs such as these, in order to further understand how metabolite conversion and intermediate accumulation leads to *in vivo* metabolic burden and whether this matches the (predicted – measured normalised capacity) calculation.

